# Functional and morphological alterations of the light perception circuits in postmortem retina from donors at different stages of Alzheimer’s disease

**DOI:** 10.1101/2025.02.18.638646

**Authors:** Nicolae Sanda, Dan Milea, Eniko Kovari, Yixin Tong, Talayeh Brügger, Jens Hannibal, Kristof Levente Egervari, Johannes Alexander Lobrinus, Emmanuel Carrera, Gabriele Thumann, Sonja Kleinlogel, Antoine Adamantidis, Ludovic S. Mure

## Abstract

Disruption of sleep and circadian rhythms is one of the earliest symptoms of Alzheimer’s disease (AD). Circadian entrainment and modulation of alertness are non-visual responses to light driven by intrinsically photosensitive retinal ganglion cells (ipRGCs).

To explore structural and functional changes of ipRGCs and ipRGC circuits in AD, we analyzed the retinas and brains of 13 elderly patients ranging from normal cognition to AD and performed *ex vivo* extracellular electrophysiological recordings on freshly harvested retinas.

While rods and cones were moderately impaired, there was a severe loss of ipRGCs in AD donors. Importantly, the remaining ipRGCs exhibited morphological alterations, hyperexcitability, and were not able to sustain high levels of activation. These changes may be ipRGC subtype-specific and correlated with disease progression.

Altered ipRGC circuits and function could contribute to the disruption of sleep and circadian rhythms reported in AD patients. Measuring ipRGC-dependent responses to light could be a promising way to predict or monitor pathological changes in the brain.

## Introduction

Alzheimer’s disease (AD) is a devastating neurodegenerative disorder and the most common cause of dementia. characterized by progressive cognitive decline, memory loss, and impaired daily functioning^1^. AD neuropathological features, such as the presence of amyloid-beta plaques and tau protein-containing tangles, play a central role in disease progression. A definitive and unambiguous diagnosis is difficult to establish, and by the time it is obtained, the disease has usually progressed over decades, causing significant neurodegeneration. It is estimated that dementia affects more than 55 million patients, however its prevalence continues to rise as the population ages, and it is projected to double by 2050^2^. Understanding the underlying disease mechanisms and identifying potential early biomarkers is urgently needed to allow timely diagnosis and intervention.

One of the earliest symptoms in AD is a disruption of sleep and circadian rhythms. Individuals with Alzheimer’s often experience increasingly fragmented sleep and an altered core body temperature rhythm^3–6^. There is now abundant evidence that disruption of the daily rhythms can occur years before the emergence of classical symptoms, such as memory loss in AD^7,8^. Excessive daytime sleepiness and sleep behavior disorders are associated with an increased risk of developing AD and dementia suggesting that dysregulation of sleep may be important in the early pathogenesis of neurodegeneration^9–12^. These behavioral disruptions compromise not only the quality of life of the patient, but they also cause physical and psychological fatigue in the caregiver, often resulting in institutional placement.

The regulation of non-visual responses to light, including the synchronization of the circadian clock to the day-night cycle, as well as the regulation of sleep, alertness, mood and the pupillary light reflex (PLR), is primarily mediated by a class of intrinsically photosensitive retinal ganglion cells (ipRGCs) that express the light-sensing protein melanopsin. Accordingly, when ipRGCs are selectively ablated or when signaling properties of melanopsin are changed, photoentrainment of the clock, acute light-modulation of sleep, and PLR to light are abolished or maladapted^13–16^. Interestingly, a recent histological study has shown a specific decline in the number of ipRGCs in the retina of AD patients^17^. Beta amyloid accumulation was also observed inside and around ipRGCs suggesting that the response properties of ipRGCs from AD patients may be altered^17^. In mouse models of AD, ipRGC anomalies have also been reported^18,19^. While these observations suggest that light perception circuits may be damaged in Alzheimer’s disease, this has never been investigated further in patients.

The aim of this study was to evaluate the alteration of light perception in AD patients compared to normal individuals, using a comprehensive exploration in eye-brain donors. Specifically, we examined the histological and functional changes in ipRGC circuits in the retina and brain of individuals at different stages of Alzheimer’s disease. We found that ipRGC-mediated photoreception was severely altered in AD; a fraction of ipRGCs underwent degeneration, while the remaining cells exhibited morphological alterations, hyperexcitability, and an inability to sustain high levels of activation. These changes may have been subtype-specific and correlated with disease progression. Although our findings are limited by the small size of our sample, they emphasize the importance of further investigating ipRGCs in order to manage symptoms associated with ipRGC-mediated responses to light, to develop diagnostic tools, to design novel therapeutic strategies and, more generally, to unravel early the pathophysiological mechanisms leading to neuronal degeneration in AD.

## Results

To assess the impact of AD on light perception, we conducted an exploratory, multi-angle study on 13 eye and brain donors, deceased at the Geneva Stroke Center (Geneva, Switzerland, **FigS1A**). The demographic characteristics of the cohort are presented in **Table 1**. Ages ranged from 69 to 103 years (81.31 ± 3.011). There were 9 males (mean age 79.44 ± 3.17 years; 69 to 97 years) and 4 females (mean age 85.50 ± 7.01 years; 70 to 103 years, p=0.49, Mann Whitney test). The cause of death was brain infarct and/or brain hemorrhage (**Table1**). Among the included donors, two showed ante-mortem signs of dementia (donors 3 and 5), which, together with their post-mortem neuropathology, support a diagnosis of Alzheimer’s disease and are therefore referred to in this manuscript as AD donors, as opposed to donors without known cognitive impairments (ND, non-demented, donors). The neuropathological severity of tau lesions and amyloid deposition were assessed in the anterior hippocampus, entorhinal, transentorhinal, and inferior temporal cortices using the Braak staging for neurofibrillary tangles (NFT) and the Thal phases for senile plaques (SP) (**FigS1B-D)**^20,21^. Thal phases ranged from 0 to 4 and Braak scores ranged from I to VI. The two AD patients scored 3 and 4 and V and VI respectively **(Table1**). While Thal phases were correlated with the age of the donors, Braak stages were not (**FigS1E**).

**Table 1:** Donor’s demographic and pathology. Amyloid and tau pathologies were evaluated with Thal phases and Braak stages respectively.

|  | *Age (years)* | *Sex* | *Delay to Brain collection (hours)* | *Delay to Eyes collection (hours)* | *Cause of death* | *Thal $\beta$ Amyloid (A)* | *Braak NFT (B)* | *CERAD (C)* | *CAA* | *Brain histology* | *Retina histology* | *Retina function* | *Figure symbols* |
| --- | --- | --- | --- | --- | --- | --- | --- | --- | --- | --- | --- | --- | --- |
| *D1* | 71 | M | 6 | 2 | Brain infarct, hemorrhage | 0 (0) | II (1) | 0 | + | X | x | x | ● |
| *D2* | 80 | F | 25 | 2 | Brain infarct | 2 (1) | III (2) | 2 | - | X | x | x | ■ |
| *D3* | 77 | M | 13 | 2 | Brain hemorrhage | 4 (3) | VI (3) | 3 | + | X | x | x | ▲ |
| *D4* | 85 | M | 34 | 2 | Brain hemorrhage | 2 (3) | I (1) | 0 | + | X |  |  |  |
| *D5* | 89 | F | 114 | 1.5 | Brain infarct | 3 (2) | V (3) | 2 | + | X | x | x | ▼ |
| *D6* | 97 | M | 27 | 9 | Brain infarct | 4 (3) | II (1) | 0 | + | X |  |  |  |
| *D7* | 103 | F | 3 | 1 | Brain infarct | 1 (1) | II (1) | 0 | - | X |  |  |  |
| *D8* | 74 | M | 11 | 0.5 | Brain infarct | 1 (1) | II (1) | 0 | - | X | x | x | ○ |
| *D9* | 74 | M | 46 | 2 | Brain infarct | 0 (0) | I (1) | 0 | - | X |  |  |  |
| *D10* | 70 | F | 82 | 82 | Brain hemorrhage | 0 (0) | II (1) | 0 | - | X |  |  |  |
| *D11* | 77 | M | 41 | 2 | Brain hemorrhage | 2(1) | II (1) | 0 | - | X | x |  | ◆ |
| *D12* | 69 | M | 20 | 2 | Brain infarct | 0 (0) | I (1) | 0 | - | X |  |  |  |
| *D13* | 91 | M | 18 | 2 | Brain hemorrhage | 2 (1) | 0 (0) | 0 | - | X | x | x | □ |

To quantify AD hallmarks along some of the brain circuits implicated in the regulation of visual and non-visual responses to light, we also examined tau pathology and amyloid deposition in 13 brain regions implicated in vision (LGN, SC, visual cortex), circadian rhythms (SCN), alertness and sleep regulation (POA, pons nuclei), mood (AMY, LHB/pLHB, ACC), and pupillary reflex to light (EWN) in 13 donors (**Fig1A**). Uniform brain sampling was performed by a neuropathologist and SP and NFT were scored on a 0 to 3 scale by trained pathologists (0=none, 1=sparse, 2=moderate, 3=abundant) (**Fig1B**). We found aggregates in most of the regions examined. While deposits of Amyloid beta (Aβ) were more abundant as the amyloid pathology progresses, hyperphosphorylated tau (p-tau) burden did not follow this pattern and its presence in light perception circuits appeared uncorrelated with Braak stages (**Fig1C, FigS2A**). Aβ burden in the VC and Acc as well as in all retino-recipient nuclei except the LGN correlated with the Thal phase of the respective donor (**Fig1C**).

**Figure 1:**
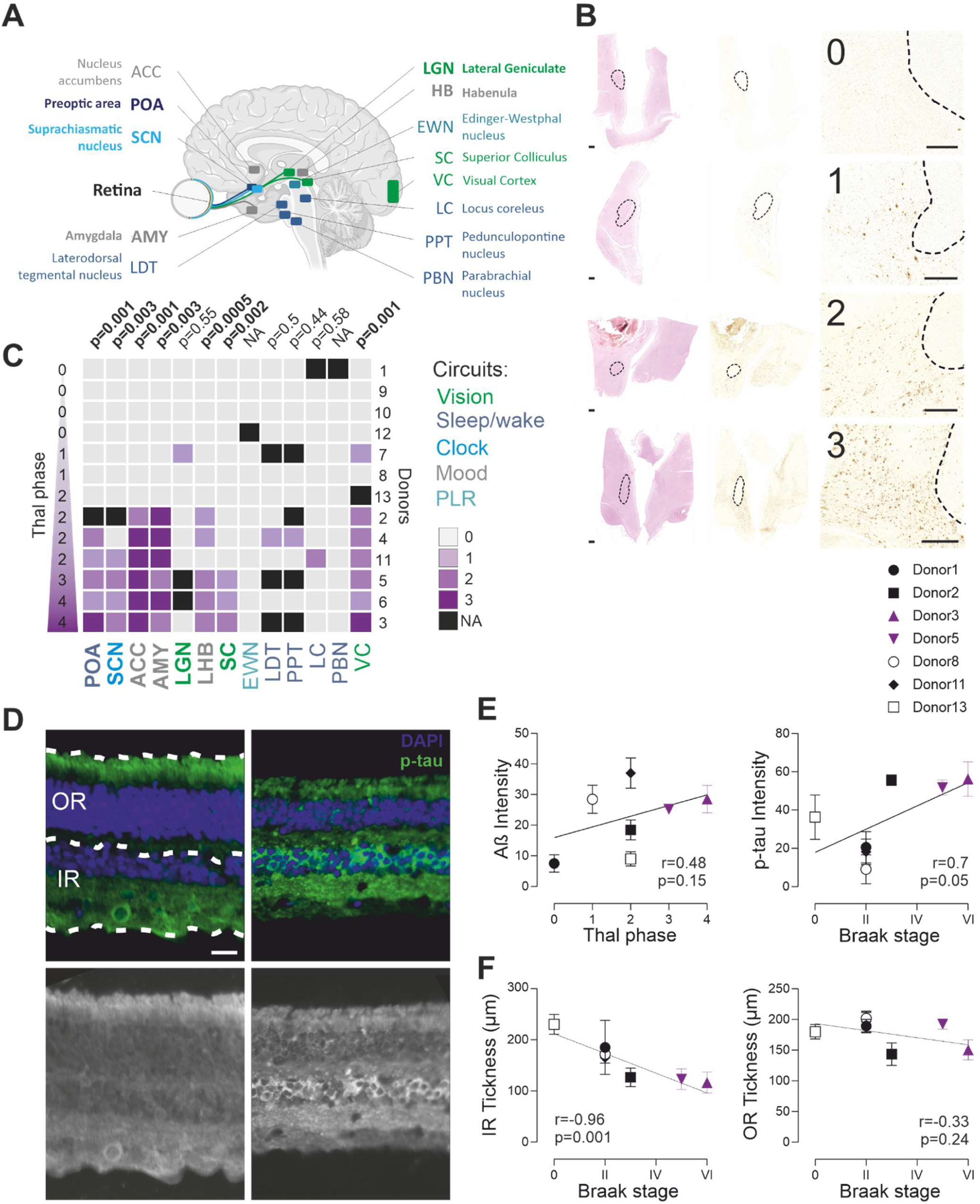
AD hallmarks are present along the circuit of light perception. (**A**) Several brain nuclei contributing to the main light-regulated function (vision, entrainment of the circadian clock, PLR, sleep/alertness, mood) were collected. (**B**) Hypothalamic region including POA from 4 donors exhibiting increasing levels of Aβ deposits (scored 0 to 3). *Left, middle, and right columns,* H&E staining, Aβ staining, and related enlargement respectively (*dotted lines*, fornix; scale bars, 1mm). (**C**) Aβ deposits scores for each brain nuclei in each donor ranked as a function of their Thal phase. (**D**) Fluorescence micrographs of retinal cross-sections from donor 7 (ND, left column) and donor 3 (AD, right column) with Thal phases 2 and 5 and Braak scores II and VI respectively. On the upper left panel, dotted lines materialized the definition of the Outer and Inner retinas (OR and IR). (*Upper row*) Tissues were immunolabeled for p-tau (AT8, green), and DAPI+-nuclei (blue;). Scale bar: 50 µm. (*Lower row*) Greyscale micrographs the same individuals immunolabeled with p-tau (Scale bar: 50 µm). Scatter plots of the thickness of the donors’ IR (**E**) and OR (**F**) as a function of their Braak score (n= 3 to 9 sections measured, Spearman correlation coefficient and p value). Scatter plots of the Aβ (**G**) and p-tau (**H**) burdens measured in each donors as a function of the Thal phase and Braak scores of the donors respectively (n=3 to 5 for Aβ and n=3 for Tau per donor, Spearman correlation coefficient and p value).

To evaluate the correlation between the brain and retina pathologies, we performed immunostaining of Aβ and Tau in the retina of the donors. We manually segmented the outer retina (OR: photoreceptor (PR) and outer plexiform layers (OPL)) and inner retina (IR: inner nuclear (INL), inner plexiform (IPL), and retinal ganglion cell (RGC) layers) (**Fig1D**) and determined the layer distribution of p-tau and Aβ burdens in retinal cross sections. Representative microscopic images show an accumulation of p-tau in patients that correlates with the Braak stage (**Fig1E**). Immunostaining of p-tau revealed diffuse staining in the OPL and intense staining in the INL and IPL, including probable intracellular staining in the INL. Aβ deposits were identified in the inner retina, in particular in the innermost RGC and retinal nerve fiber layers. Aβ42 burden (**Fig1E, FigS2B**) also increases with the Thal phase of the donor. Finally, we also measured the thickness of the segmented IR and OR as a proxy of neurodegeneration. The thickness of IR layer decreased with AD and was negatively correlated with the increase of Thal phases and Braak stages (**Fig1F, FigS2C**).

The presence in the retina of both Aβ and Tau aggregates and degeneration (**Fig1**) suggested a possible alteration of the light information sent to the brain. To evaluate the function of the rods, cones, and ipRGCs, we recorded responses to light from a donor’s retina preparation using multi-electrode array (MEA)^22,23^. Briefly, the eyes of the donors were collected within 2 hours postmortem (**Table1**), immediately preserved in oxygenated Ames ‘medium placed in a light-proof container, transported to the lab where circular (3mm diameter) patches of retina were dissected under dim red light, placed on MEA, and allowed to recover in darkness for at least 1 hour.

First, to evaluate photoreceptor function, we recorded micro-electroretinograms (mERGs)^24^ on retinal patches from five donors (**Table1**), including the two AD donors in response to sequences of 10ms light flashes of increasing irradiance (470nm, 6.10^9^ to 2.10^13^ photons/cm^2^/s) (**FigS3**). mERG did not display major differences in response profiles between AD and age-matched control donors (**FigS3A**). A-waves were detected in all donors at all light irradiances used while b-waves were only clearly detected at the highest light stimulation but in all donors too^23^. While the amplitude of the a-wave from the AD donors was the smallest, it was not different from that of the other donors. Similarly, the b wave amplitude and the latencies of both waves did not differ between donors (**FigS3B, C**).

Then, to evaluate ipRGCs’ function, we performed extracellular electrophysiological recordings of freshly harvested retina from 6 donors including the two AD donors (**Table 1**). For this purpose, the recording medium was supplemented with synaptic blockers to block rod and cone input to RGCs and capture ipRGCs intrinsic, melanopsin-initiated signal. In response to 30s pulses of monochromatic blue light (470 nm), photoresponsive cells were found in retinas from each of the six donors (**Fig2A**). However, the number of ipRGCs varied from one donor to the other. For each retina sample, we identified photoresponsive cells at an average density ranging from <1 to ∼8 cells/mm^2^ (**Fig2B**). The number of light responsive cells was negatively correlated with the Braak stage but not with the Thal phase, the age of the donor, or the delay of collection of the eye (**Fig2B and FigS4A**). Grouped analysis reveals a two-fold decrease in ipRGCs density in retina pieces from AD donors (**FigS4B**).

**Figure 2:**
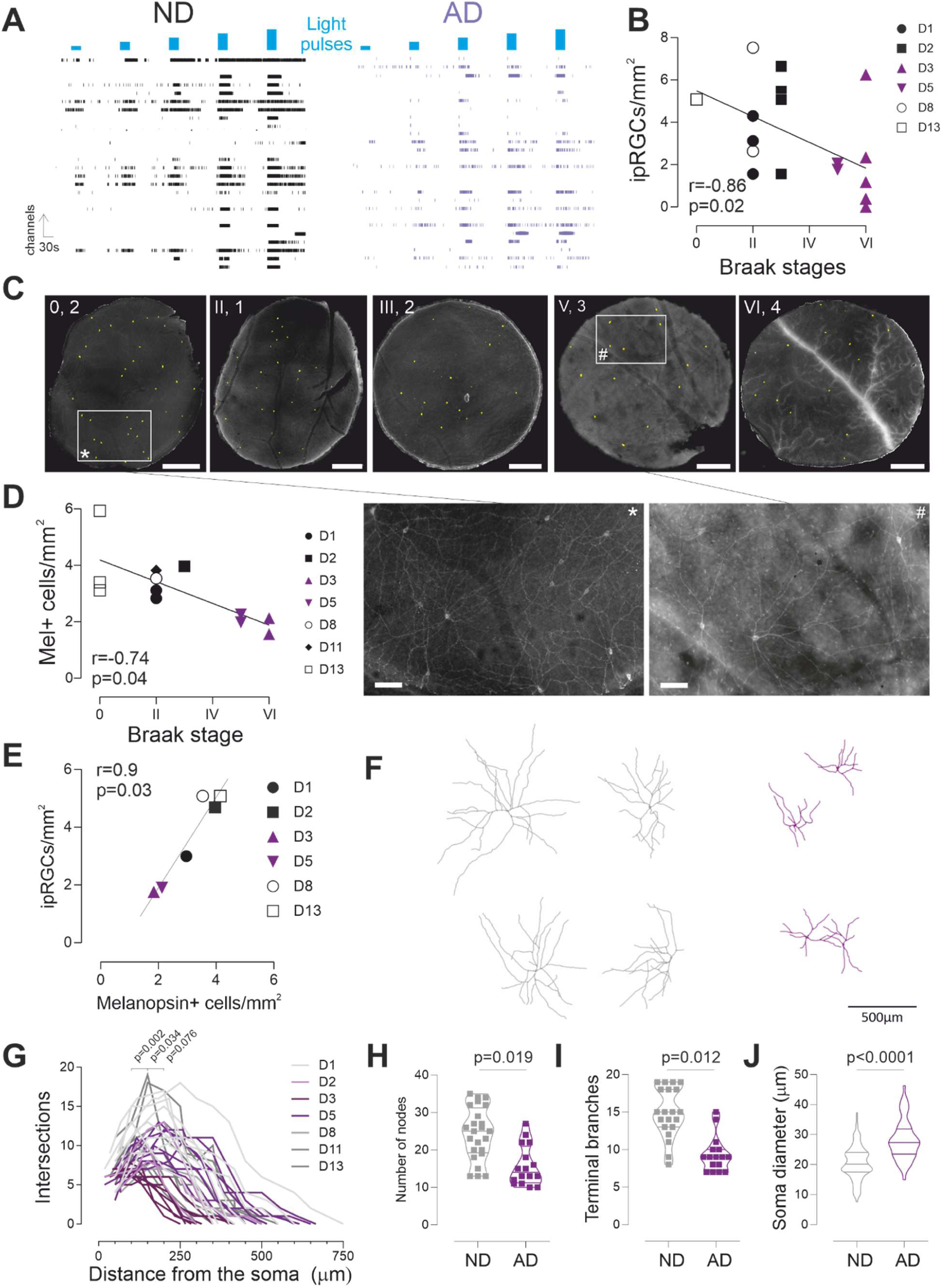
ipRGCs decline in AD. **(A)** Representative ipRGC spike trains to a 30-s light pulses of increasing irradiance (∼10^10^ to 10^15^ photons/cm^2^/s, 470 nm) from AD and age-matched ND donors. (**B**) ipRGCs density in pieces of retina from each donor as a function of their Braak score (n=1 to 5 retina pieces per donor). (**C**, *Upper row*) Microphotographs of patches of retina immunostained against melanopsin (scale bar 500µm, *upper right corner*, Braak stage and Thal phase of the donors). (*Lower row*) Enlargements from the first (*) and fourth (#) patch microphotographs (scale bar 100µm). The mel+ cells density in each donor correlates with their Braak score (**D**, n=1 to 2 retina pieces per donor) and with the density of ipRGCs reported in the same donors (Spearman correlation, **E**). (**F**) Examples of dendritic shrinkage, mel+ cells from AD (Donor3, *purple*) and age-matched (Donor1, *grey*) donors (scale bar 500µm). (**G**) Scholl analysis (Linear Mixed-effect model (LMM), interaction F(14,555)=3.7263, p<0.001, FDR-corrected post hoc contrast p-values on the figure), (**H**) number of nodes, (**I**) number of terminal branches, and (**J**) soma diameter from AD (n=8 cells per donor, 2 donors, *purple*) and age-matched (n= 4 to 8 cells per donor, 4 donors, *grey*) donors were different (LMM, F(1,38)=5.99 and F(1,38)=6.98, H and I respectively, F(1,199)=36.522, J).

To confirm the ipRGC loss in AD patients, we performed immunostaining against melanopsin as previously described^25^ and we counted melanopsin-positive (mel+) cells in 3 mm diameter retinal patches (**Fig2C**). We found about two times decreased density of mel+ in the retina of AD donors (**FigS4B**). When examined individually, mel+ density correlated with the patient Braak stage and ipRGCs density, but not with the Thal phase (**Fig2D, E and FigS4C**). Melanopsin staining revealed that not only the number, but also the morphology of mel+ cells is affected by the disease. In each retina sample, a blinded investigator used semi-automated tracing to reconstruct the dendritic arborization of 4 mel+ cells per retina sample (**Fig2F**). Scholl analysis revealed that while the extend radius of the ipRGCs dendritic tree was not different between donors, its number of intersections midway between the soma and the extremities was decreased in AD donors (**Fig2G, FigS4D**). Accordingly, both number of nodes and terminal branches were lower in the AD patients (**Fig2H-I, FigS4D**). These results suggest a decrease in dendritic tree complexity in AD. Interestingly, AD ipRGCs had larger soma (**Fig2J, FigS4D**). Altogether, these major morphological alterations strongly suggest that the function of the surviving ipRGCs is also altered.

To evaluate potential functional alterations in the remaining ipRGCs from AD donors, we characterized their response properties. We stimulated the retinas for 30 s at increasing irradiances of monochromatic blue light (470 nm, 2.10^12^ to 1.10^15^ photons/cm^2^/s). Overall, responses to light were slow, sustained, and outlasted the stimulus by several seconds after the light was switched off, the sensitivity was low as previously described^22^ (**Fig2A**). Response properties followed similar dynamics with increasing stimulation irradiance; latency decreased while the duration and number of spikes of the response increased (**Fig3A**). Interestingly, responses from donors at more advanced stages of the disease appeared stronger at lower light intensities reflecting an increased sensitivity or excitability with the progression of the disease (**Fig3B,C**). Conversely, in these donors, responses to the highest intensity of light were diminished or even sometimes abolished, suggesting a failure to sustain responses at advanced stages of AD (**Fig3B,C**).

**Figure 3:**
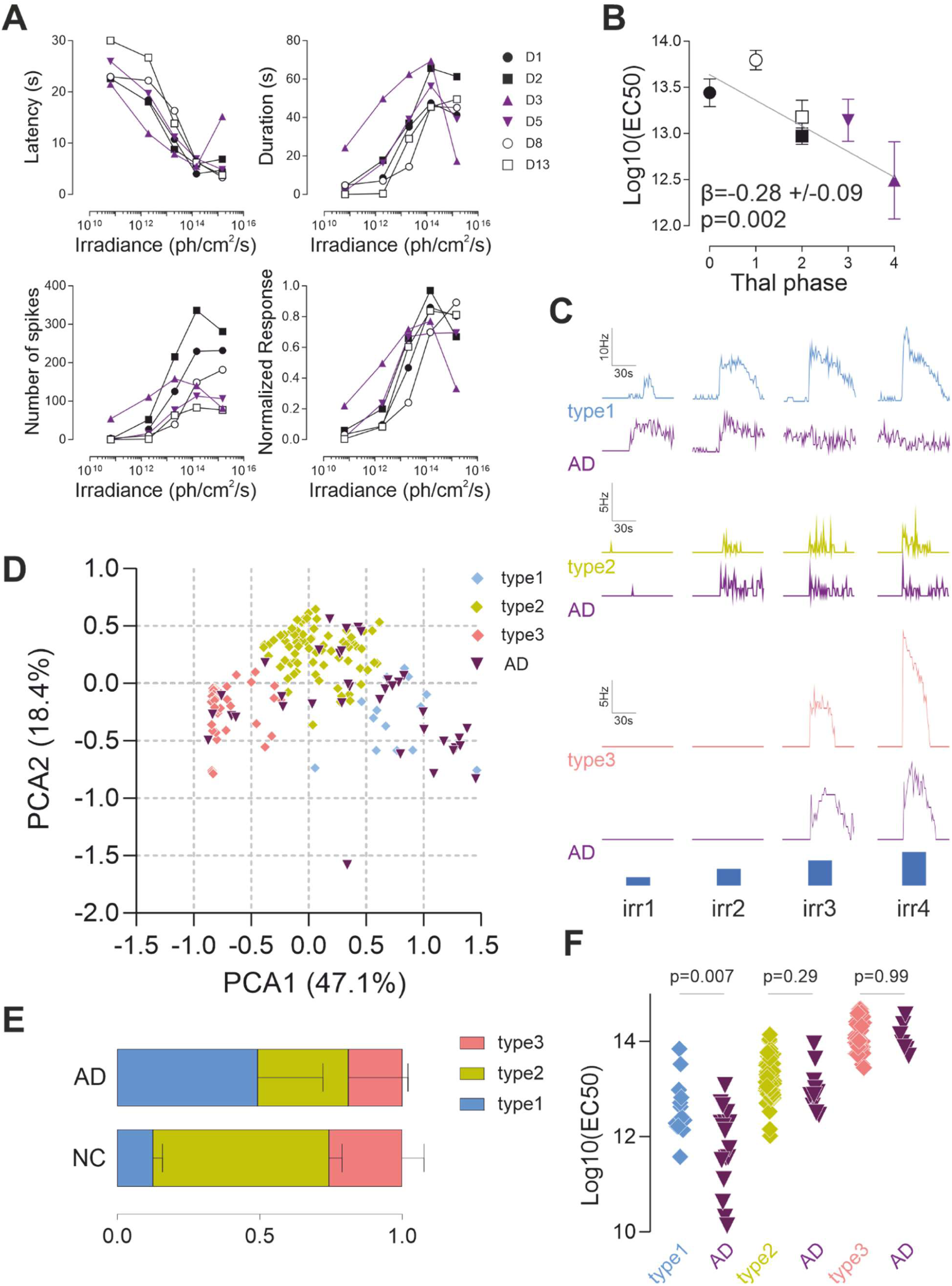
ipRGC function is altered in AD. (**A**) Average ipRGCs’response latency, duration, number of spikes, and normalized response in each donor (n=28, 15, 18, 19, 77, and 13 for donors 1, 2, 3, 5, 8, and 13 respectively, where n is the number of cells) to a sequence of 30s light pulses of increasing irradiance (∼10^10^ to 10^15^ photons/cm^2^/s, 470 nm). (**B**) Average ipRGC sensitivity to light as a function of Thal phase (LMM, slope β +/−SE). (**C**) Representative responses from types 1, 2, and 3 ipRGCs to increasing irradiance light pulses (30 s, 470 nm, irradiances (irr) 1, 2, 3, and 4: 2×10^12^, 2×10^13^, 1.5×10^14^, and 1.5×10^15^ photons/cm^2^/s, respectively). Blue bars indicate light pulses. (**D**) Principal components of the ipRGCs’ response parameters (sensitivity, latency, and duration; n =133 and n=37, four control and two AD donors (*purple*) respectively). (**E**) Distribution of ipRGC subtypes in ND and AD donors. (**F**) ipRGC sensitivity by subtype in ND and AD donors (n=14, 77, and 45, type1, 2, and 3 cells and n=18, 12, and 7 type1, 2, and 3 cells respectively, LMM F(1,30)=8.43 type 1, F(1,87)=1.09 type 2, F(1,50)= 2.2.10^−6^, type 3).

Several morphological and functional ipRGC subtypes have been described in the human retina^22,25–28^. To assess whether the alterations reported above reflect alteration and/or degeneration of specific subtype(s), we performed principal components analysis (PCA) on the response parameters of each ipRGCs recorded. Data separated into at least three main clusters (**Fig3D**) whose response to light properties and time course closely resemble those from previously described ipRGC subtypes^22^ (**Fig3C,D,**). The three subtypes were found in all donors, but their distribution was changed in the AD donors. Type 1 accounted for half of the ipRGCs in AD donors while type 2 and 3 ipRGCs fraction was strongly decreased (**Fig3E**). When examined individually, type 1 fraction correlates with the Braak stage of the donor (**FigS4E**). This result suggests that type 1 cells may be more resilient than type 2 and 3. As type 1 ipRGCs are the subtype that is most sensitive to light, the increased excitability observed could be due to this change of distribution. To test this hypothesis, we examined the sensitivity to light of each subtype. We found that type 1 ipRGC sensitivity was increased in AD donors (**Fig3F**). Overall, all ipRGC subtypes are affected and the increased excitability observed in AD may rely on both a shift in subtypes distribution toward type 1, the most sensitive ipRGCs, and an increase in their sensitivity.

## Discussion

In this prospective study, we examined several circuits of light perception in a cohort of elderly adult donors at various stages of AD pathological changes. Using a comprehensive suite of measurements, we provide the first picture of how AD may impact ipRGCs abundance, function, and brain targets. We found that the light signals conveyed by ipRGCs to the brain are largely altered with AD, both quantitatively and qualitatively. These changes seem to correlate with the progression of the disease and may affect subtypes selectively. Importantly, we highlight the potential of ipRGCs as both biomarkers for AD and models to study the mechanisms of neuronal degeneration in humans.

In the retina, our results are in agreement with the presence and distribution of AD hallmarks and their correlation with the neuropathological changes observed in the brain, reported early and in extensive postmortem retinal histological studies (^29–33^(for review see^34^)). We also confirmed the loss of ipRGCs observed in AD patients^17^. We extended these findings by measuring *ex vivo* the function of the different photoreceptors. Both outer and inner retina photoreceptors seem to malfunction with AD but, while rods/cones response alteration may be limited to small decreases in amplitude at specific light intensity levels, ipRGCs injury is massive.

AD has been associated with various forms of visual dysfunction, including abnormal visual fields, reduced color and contrast vision which can be, at least in part, attributed to retinal and optic nerve damage^35–38^. Retinal abnormalities can be identified with electroretinogram (ERG) in AD ^39–43^. Here, while we found a decrease in IR thickness, the mERG a and b waves did not show significant differences between AD and ND patients in response to our stimulation protocol. Because IR photoreceptor function declines rapidly with hypoxia^23^, some differences between donors may have been masked by the postmortem functional decline in all donors.

ipRGCs are the main contributors and in some cases the exclusive conduits of light regulation to non-visual light responses^13,14^. Accordingly, they project to more than a dozen of brain regions implicated in visual and non-visual responses to light, including the nuclei investigated here: SCN, POA, HB/pHb, AMY, LGN, and SC^44,45^. We found twice less ipRGCs in AD donors, and the surviving ones displayed abnormal morphologies and responses to light that may precede degeneration. In particular, the increase in soma size resembles the phenomenon called neuronal ballooning that is thought to result from deficient axonal trafficking and subsequent accumulation of cargos at the initial segment^46–48^. Then, the increased excitability is reminiscent of cerebral neurons in which an increased excitability is observable at the earlier stages of the disease before there is pronounced cell loss and hypoactivity^49–51^. The initial loss of ipRGCs may be biased toward types 2 and 3, with type 1 ipRGC appearing more resilient. Of note, while the response profiles of ipRGC subtypes in AD donors seem to remain similar to their counterparts from control donors (**Fig3C**), this result would need confirmation as the properties on which the classification is based are altered and thus could lead to misclassification. M1 ipRGCs, the morphological equivalent of functional type1 ipRGCs, had already been shown to demonstrate increased survival to mechanical and metabolic stresses^52^. However, the morphological and functional alterations observed in AD donor remaining ipRGCs suggest that type1 ipRGCs probably also degenerate. If confirmed, these results would suggest that some ipRGC-dependent responses to light may be impacted earlier than others and could constitute stage-specific markers. Overall, these alterations are consistent with and could contribute to the alterations of ipRGC-regulated functions, such as circadian rhythm, sleep/alertness, and mood, experienced by AD patients. Interestingly, large-scale population studies, such as those from the UK Biobank, provide indirect support to our results by reporting associations between light signaling, sleep, and neurodegenerative outcomes^53–55^.

As PLR is thought to rely on both M1 and M2 ipRGCs^56^, our results predict that pupillary responses to light should also be affected. Pupillometry is a scalable, cost effective, non-invasive assay that could easily be used in clinical settings. A few groups have started to explore as a biomarker of AD in pre-symptomatic cases^57,58^ and early AD cases^59,60^. However, the results have been mixed. This may reflect the diversity of light stimuli and protocols^61,62^ and emphasizes the need to determine the spectral, intensity, and/or temporal domains where ipRGCs malfunction and PLR is impacted. For example, our results suggest that injured ipRGCs will fail to sustain responses to long stimulations at high irradiance.

The retina, an extension of the central nervous system, offers a unique opportunity to study AD pathology outside of the brain. Over the past decades, accumulating evidence has shed light on the presence of AD-related histological changes in the retina, suggesting the potential of different retinal imaging modalities as a non-invasive method to detect and study the disease (for review see^34^). Our results suggest that the retinal functions (light perception and transmission to the brain) are also altered and add ipRGC-dependent responses to light as potential biomarkers of AD. In AD donors, surviving ipRGCs seem to recapitulate several phenotypes observed in brain neurons: hyperexcitability and ballooning. This, together with the fact that they are more accessible and have clear cellular and behavioral readouts, makes ipRGCs a promising model to study the mechanisms of neuronal degeneration directly in humans. Elucidating the mechanisms underlying ipRGC vulnerability to AD could lead to the discovery of therapeutic targets to improve patients’ quality of life and potentially improve the overall disease trajectory.

### Limitations

Our study has a number of limitations. First, our sample size is small because of the challenges presented in obtaining and collecting postmortem eye and brain tissues from well-characterized participants meeting the criteria of inclusion. However, we partially compensated for this limitation by using in-depth objective physiopathological and functional measurements from each donor. Secondly, our cohort has a greater proportion of male subjects, leading to potential sex-related bias. Our study population was composed primarily of self-reported White and thus may not be representative of other populations. Third, because our focus was on physiology, the pieces of retina were collected in the same central region, irrespective of the orientation. While this strategy may have caused some variability as ipRGC density varies between people^28^, ipRGC density and distribution is relatively homogenous in this region within subject^25^. It may also have left apart more important alterations in the peripheral retina^17,29,63,64^. Finally, our study is primarily correlational and, therefore, caution must be taken before making cause-and-effect inferences.

Despite these limitations, the present study provides a first glimpse of the pathophysiological mechanisms of neurodegeneration at work in humans. Quantitative studies in larger cohorts and animal models of AD will be necessary to confirm these results and mechanistically dissect the processes underlying ipRGC vulnerability to AD. This work has the potential to influence both AD research and clinical practice by highlighting the retina as a window into brain pathology.

## Methods

### Donors

The donors were patients of both sexes from the Geneva Stroke Center who had died following a vascular event. Aside from the stroke, they were either cognitively healthy, diagnosed with Alzheimer’s disease (AD), or showed signs of neurodegeneration. Patients were prospectively included in the study after obtaining informed consent from family members or legal healthcare proxies. In a second phase, following confirmation of death, the patients were promptly transported to the hospital morgue, where enucleation was performed immediately, day or night, by a trained physician (NS). After collection, the eyes were immediately transferred to the laboratory in a light-proof, thermally insulated container filled with continuously oxygenated Ames’ medium, as previously described ^22,23^. Only patients for whom enucleation was completed within two hours post-mortem were included in the study (Table1), as photoreceptor function can be reliably recovered and recorded within this time window ^22,23^.

### Eye sampling procedure

Bilateral enucleation was performed in all eligible patients following a standard surgical procedure. After insertion of a lid speculum, a conjunctival peritomy was performed using a #15 blade, followed by dissection of Tenon’s capsule. The extraocular muscles were identified, isolated, and transected, facilitating identification and isolation of the optic nerve from the surrounding structures. Using microscissors, the optic nerve was then sharply transsected, as far as possible from its bulbar insertion to the ocular globe, closest possible to the orbital apex. After being freed from its orbital attachments, the eyeballs were removed from the orbital cavity. A 360-degree concentric limbal incision allowed the removal of cornea, iris, and the lens. Following this procedure, eyeballs were transported to the laboratory in a custom-designed container under continuous oxygenation as described previously^22,23^.

Exclusion criteria were the presence of known significant posterior pole ocular pathology such as age-related macular degeneration, glaucoma, or other retinopathies or optic neuropathies as well as know systemic pathology like diabetes.

### Brain histopathology

The tissue blocks were embedded in paraffin and 12 μm-thick adjacent sections were stained with hematoxylin-eosin or with antibodies against tau (AT8; Pierce Biotechnology, Rockford, IL, USA, 1:1,000, that recognizes the Ser202 and Thr205 epitopes) and Aβ (4G8, Signet Laboratories, Dedham, MA, USA, 1:2,000, that recognizes amino acids 18–22 of the Aβ protein). The neuropathological assessment of Alzheimer disease pathological changes was performed following NIA-AA guidelines^65^. The severity of tau--immunoreactive lesions, amyloid deposition and neuritic plaques was determined using the Braak stages, Thal phases, and CERAD classification^20,21,66^.

### Retina sampling

The vitreous was removed from the eyecup under dim red light. The macula was identified according to its pigmentation and the pattern of blood vessels in the retina. Circular patches (3 mm diameter) of retina were punched out in an annular region comprised within ∼2 to 10 mm from the fovea (except in the blind spot) and from both left and right eyes. Given the number of challenges, including punching out adequate patches, vitreous removal, patches adhesion on the MEA, length of dark adaptation periods and protocol, sampling was done indifferently of the quadrant orientation. While orientation difference may be a source of variability, it seemed a reasonable compromise as this region is relatively homogenous regarding ipRGC density.

Samples used for histological experiments were taken in the same region at the end of the recording session and immediately drop in 4% paraformaldehyde (PFA).

### Retina histopathology

Retinas were fixed in 4% PFA at 4°C for 24 hours, followed by paraffin embedding. Sagittal-oriented sections of the posterior segment were sliced at 10µm thickness. These paraffin sections were treated with 70% formic acid (Roth, 5355.1) for 10 minutes at room temperature, followed by antigen retrieval in citrate buffer (pH 6.0, Merck, 100244) at 100 °C for 1 hour. Sections were then incubated in a blocking solution containing 5% normal goat serum (Sigma, D9663) and 0.05% Triton X-100 (Sigma, T9284) in Tris-buffered saline (TBS; Sigma, 94158) at room temperature for 1 hour. Then, sections were incubated overnight at 4 °C with primary antibodies: anti-Amyloid Beta (ABeta x-42, clone 12F4, mouse, Merck, 05-831-I, 1:500) or anti-Tau (AT8, p-tau Ser202/Thr205, mouse, ThermoFisher, MN1020, 1:250), diluted in 2% blocking buffer. Sections were washed with TBS containing 0.05% Triton X-100, followed by incubation with secondary antibodies (anti-mouse Alexa Fluor® 488, ThermoFisher, A11029, 1:1000) and counterstaining of nuclei with 4′,6-diamidino-2-phenylindole (DAPI; Sigma, D9542) for 3 hours at room temperature. Finally, sections were mounted with Inova Mounting Medium (Ruwag Handels AG, 508005). Images were acquired at 40x magnification using a scanning laser microscope (Nikon, Eclipse 80i). Three to six distinct sections imaged per donor. Controls were processed using identical protocols while omitting primary antibody to assess non-specific labelling. The outer and inner retinas, including respectively the photoreceptor and outer plexiform layers and the inner nuclear layer, inner plexiform layer and RGCs layer, were manually segmented using Fiji software (NIH). The average intensity of the aggregates fluorescent signal and the thickness of these regions were measured.

### Melanopsin Immunofluorescence

Immunohistochemistry was performed as described in detail previously^25,67^. The flat-mount pieces of retinas were incubated in 5% normal donkey serum for 20 min to avoid nonspecific staining. The flat-mount was incubated for 3–4 days with melanopsin primary antibody at 4°C. The patches were then washed and incubated with secondary antibodies for 2 to 3hours at room temperature. Control experiments were performed by eliminating the primary antibody, which abolished all specific staining. Experimenters blinded for the donor ID and pathology were asked to count all cells in the pieces of retina and to trace four representative cells in each patches using semi-automated tracing (pluggin SNT in Fiji).

### Multi-electrode array recordings

Patches of retina were dark incubated in oxygenated Ames solution until recording (at least 60min). Retinal patches were mounted on MEA (256 electrodes arranged in a 16 × 16 square grid, Multichannel Systems, Reutlingen, Germany), ganglion cell side contacting the electrodes, and perfused with oxygenated Ames’ medium (Sigma-Aldrich, A1420-10X1L) at 34°C. The MEA2100 amplifier was enclosed in a lightproof box located in a room kept in dark with light emitted only from the controller computer screen with red filter on the screen. ipRGC recordings were done in presence of synaptic blockers (CNQX 20µM, L-AP4 20µM, and D-AP5 50µM) and 9-cis retinaldehyde. Stock solutions of 9-cis-Retinal (R5754; Sigma–Aldrich) in ethanol (100 mM) were prepared in darkness and stored at −80°C. On the day of the experiment, an aliquot was thawed and diluted to a final concentration of 100 µM in Ames’ medium.

Data collection, and analysis were performed as previously described^16,22,68^. Briefly, the signal was acquired at 10 kHz and filtered using a low pass (<500Hz) and a high pass (>300Hz) filters for the mERGs and for the RGC multiunit spike responses respectively. Thresholds for spike detection were set at 5 times the standard deviation of the noise on each channel. Spike cutouts, consisting of 1 msec preceding and 2 msec after a suprathreshold event, along were sorted into trains of a single cell after recording using Offline Sorter (Plexon, Denton, TX). Data analysis and display were performed using Neuroexplorer (Plexon) and custom software written in MATLAB.

### Light stimulations

To elicit the mERGs, we used sequences of five 10ms monochromatic light pulses (470nm) separated by 10s of darkness at four increasing irradiances (6.10^9^, 7.10^10^, 2.10^12^, and 2.10^13^ photons/cm^2^/s). To generate ipRGC responses, we applied a protocol consisting in a sequence of four 30s light stimulations of increasing irradiance of light (7.10^10^, 2.10^12^, 2.10^13^, 1.10^14^, and 1.10^15^ photons/cm^2^/s at 470nm,) separated by 2min of darkness. All light stimuli were generated by high brightness LED, LuxeonStar 5 located ∼20cm at the vertical above the MEA. Stimuli were controlled by a homemade C+ program and synchronized to the electrophysiological recordings. The light intensities were measured with a powermeter (PM120D, Thor labs).

### ipRGC response parameters analysis

The response discharge rate was calculated as the average of the discharge rate measured during the stimulation (30s) minus the average discharge rate measure during the 30s prior to the stimulation (baseline). The response latency was defined as the delay between the start of the stimulation and the response crossing a threshold set at baseline +2*SD. The response was considered over after firing rate dropped below the baseline level for more than 3 seconds. The dose-response curves (DRCs) were obtained by plotting the response (Y, discharge rate) as a function of the irradiance of the stimulation (X, photon/cm^2^/s). DRCs were fitted by a Michaelis-Menten function of the form; Y=Rmax*X/(EC50+X) where Rmax is the maximum response and EC50 is the irradiance required to drive a 50 percent response. A few inhibitory responses were manually discarded.

### ipRGCs clustering

To distinguish functional types of ipRGCs, we performed principal component analysis (PCA) on sensitivity, duration, and latency of the response as previously described^22,69,70^. PCA allowed reducing the dimensionality of the data set; the first two principal components accounted for almost 65.5% of the variance (PC1=47.1% and PC2=18.4%). Thus, we next performed k-means clustering analysis on the first two principal components. This model produced clusters that corresponded to the initial analysis. To determine the number of ipRGC clusters, the first two principal components were fitted with Gaussian mixture models (GMMs). We produced separate GMMs for 1 to 5 ipRGCs clusters and then used the Akaike information criterion (AIC) to determine which GMM reported best the ipRGCs distribution. When we compared the 1 to 5 cluster models, the models with the smallest AIC value (best model) were the 3, 4, and 5 clusters model (AIC values for models at 1 to 5 clusters: 480.3, 386.9, 272.2, 257.5, 240.5, and 238,7 respectively). Given the limited number of cells and given the clear inflexion in AIC value at 3, we represented the 3 clusters model as the minimum number of functional subtypes, comparable to previous study^22^.

### Statistics

All data are expressed as means ± SEM unless stated otherwise. GraphPad Prism was used for all statistical analyses. Statistical tests used are stated in the figure’s legends. Differences between groups were considered statistically significant for p<0.05. Bivariate correlation analysis between histological (ipRGC/mRGC density, Thal/Braak scores, severity of brain pathology) and clinical data (age, collection delay) was performed using Spearman coefficient. Matlab (2019a) was used for k-means and hierarchical clustering, PCA, GMM, and AIC analysis (Statistics and Machine Learning Toolbox). Because of repeated measures on multiples pieces of retina per donor in some experiments, to assess the impact of the disease on measured parameters, we fitted a linear mixed-effects model (LMM) using MATLAB (fitlme). The model included disease as a fixed effect and random intercepts for donor and piece according to the formula: parameter ∼ disease (AD vs ND or Thal/Braak score) + (1 | donor) + (1 | piece). Model parameters were estimated by maximum likelihood (ML). Significance of fixed effects was assessed using F-tests with Satterthwaite approximation for degrees of freedom. F and p-values or slope (β) +/− standard error (SE) and p-values given in the respective figure legends.

### Study approval

Informed consent was obtained either from patients or from family members or legal healthcare proxies in accordance with the protocol approval obtained from the Ethics Cantonal Committee of Geneva (Project 2020-02126).

## Data availability

Code and data used to generate the figures described in this paper will be made available via Zenodo (Zenodo.org) at the time of publication, with a DOI to be included here.

## Acknowledgements

We thank Christian Dellenbach and Christian Kaiser for the light stimulation device and software, Dr. G. Benegiamo for critical comments on the manuscript, and the Donors and their families. This study was supported by the Velux Stiftung grant to SK and LSM, the Novartis Biomedical Research Foundation grant to LSM, and the Chan Zuckerberg Initiative grant to AA and LSM. JH is supported from the Danish Center for Cellular Communication.

## Contributions

NS, DM, and LSM conceptualized the study. NS and LSM designed it. NS coordinated tissue collections and harvests. NS, EK, YT, TAZ, KLE, and LSM performed the experiments. JH provided the melanopsin antibodies. EK, JH, JAL, EC, GT, SK, and AA provided scientific expertise and critical reading of the manuscript. LSM analyzed the data. LSM wrote the first draft, all authors edited and approved the manuscript.

