## Supplemental figures for "Functional and morphological alterations of the light perception circuits in postmortem retina from donors at different stages of Alzheimer’s disease"

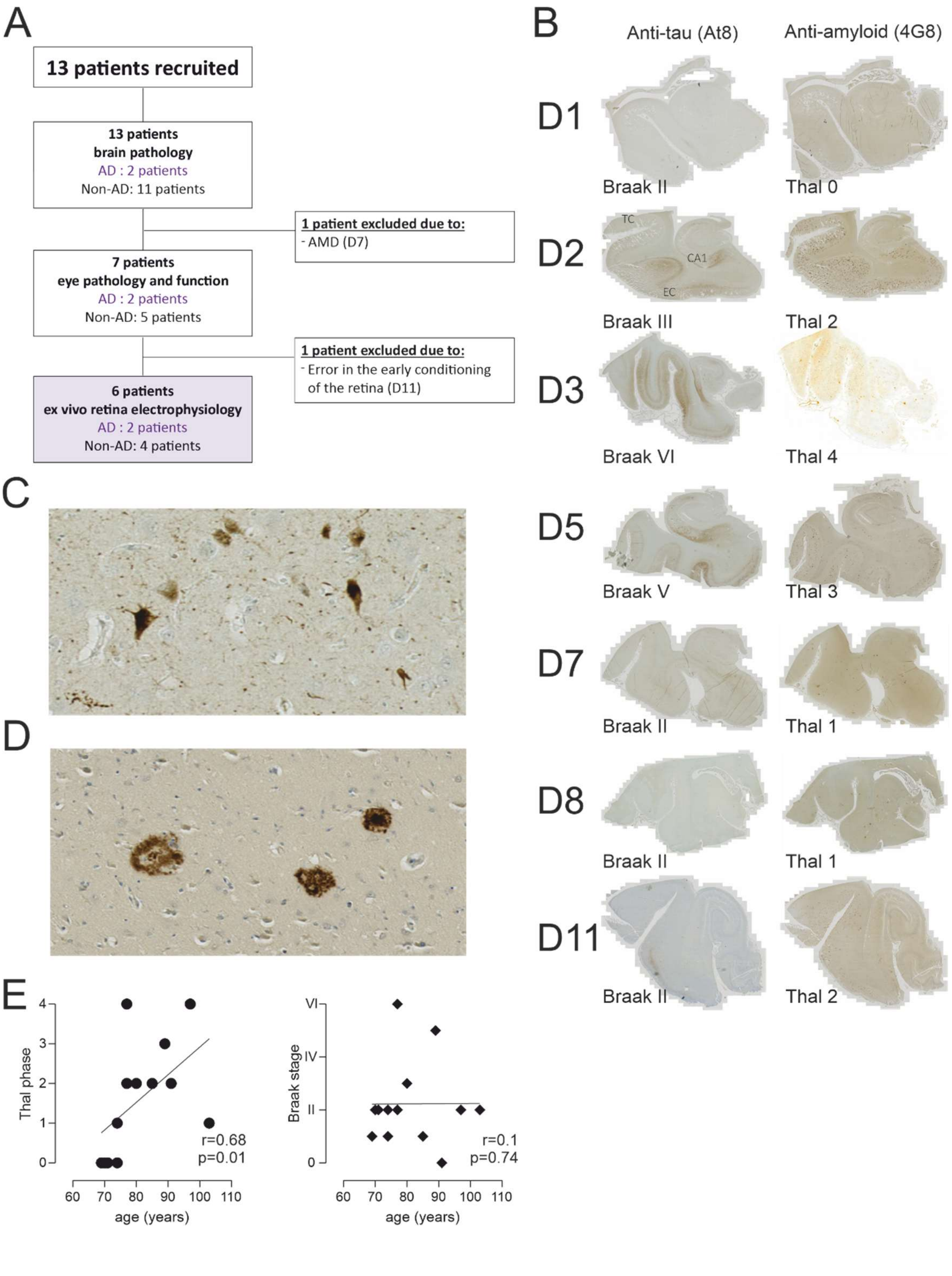


**Supplemental Figure 1: Donor's inclusion and neuropathology**. (**A**) Donor inclusion flow chart. 13 brain-eye donors were included in our protocol in 2021 and 2022 of which two had to be excluded; one due to AMD diagnostic and a second following an error in eye conditioning. 7 pair of eyes were collected within 2h and could be recorded on MEA. (**B**) NFT and Aβ deposition in the hippocampal formation and the temporal neocortex (CA1 sector of the hippocampus; EC = entorhinal cortex; TC = temporal cortex). Left column: anti-tau (AT8); right column: anti amyloid (4G8) immunostaining. High magnification of neurofibrillary tangle (**C**) and cored senile plaque (**D**) (C: AT8 tau immunohistochemistry; D: 4G8 amyloid immunohistochemistry. (**E**) Neither the Thal phase nor the Braak stage were correlated with the age of the donors (Spearman correlation).

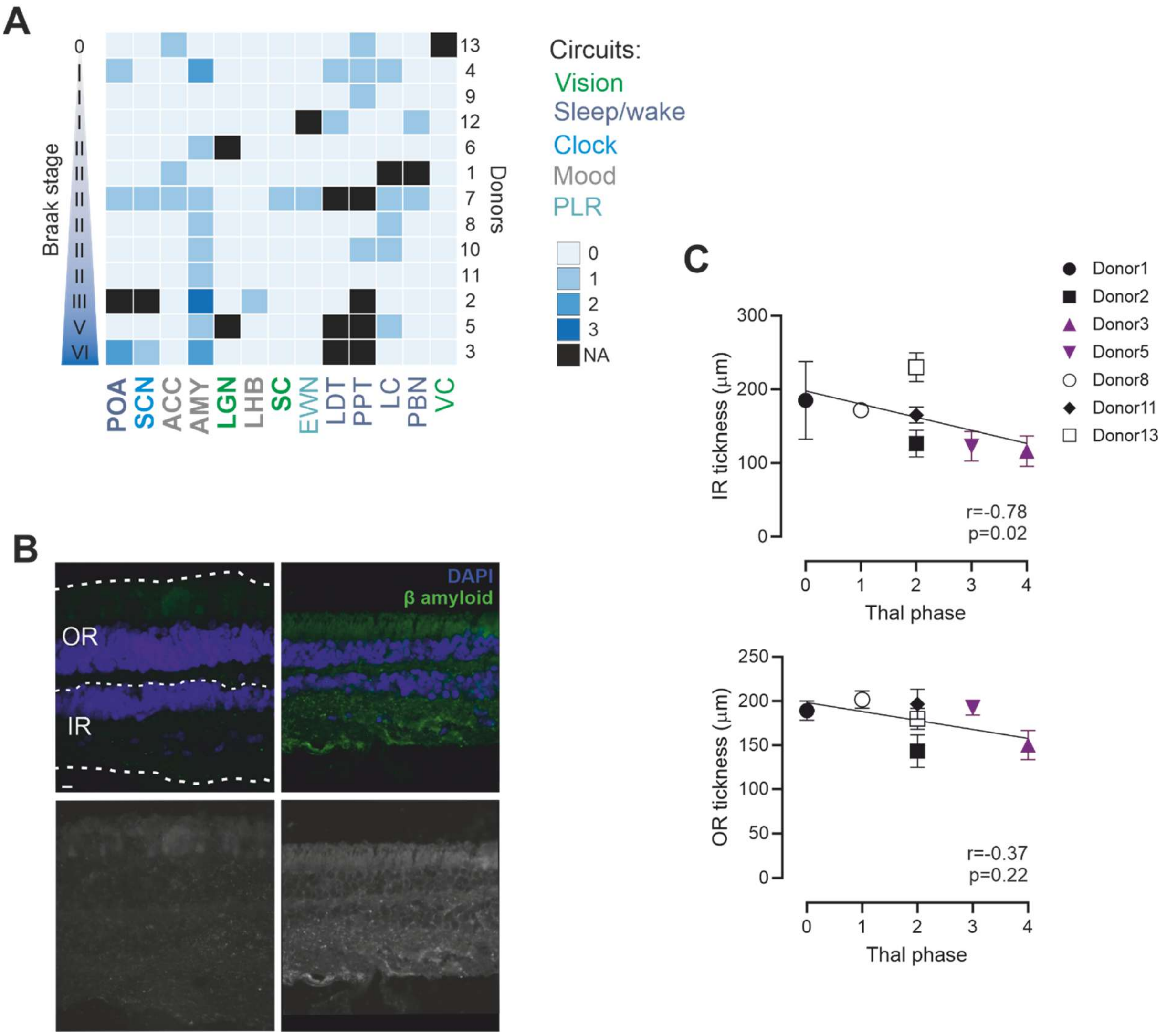


**Supplemental Figure 2: AD hallmarks in the brain and in the retina.** (**A**) Heatmap of the tau deposits scores for each brain nuclei in each donors ranked as a function of their Braak score. (**B**) Fluorescence micrographs of retinal cross-sections from donors 7 (left column) and 3 (right column) with Thal phases 2 and 5 and Braak stages II and VI respectively. On the upper left panel, dotted lines materialized the definition of the Outer and Inner retinas (OR and IR). (*Upper row*) Tissues were immunolabeled for 12F4+-Aβ42 (green), and DAPI+-nuclei (blue;). Scale bar: 50 µm. (*Lower row*) Greyscale micrographs the same individuals immunolabeled with 12F4+-Aβ42. (**C**) Correlation of outer and inner retina thickness with the Thal phase of the donors (Spearman correlation).

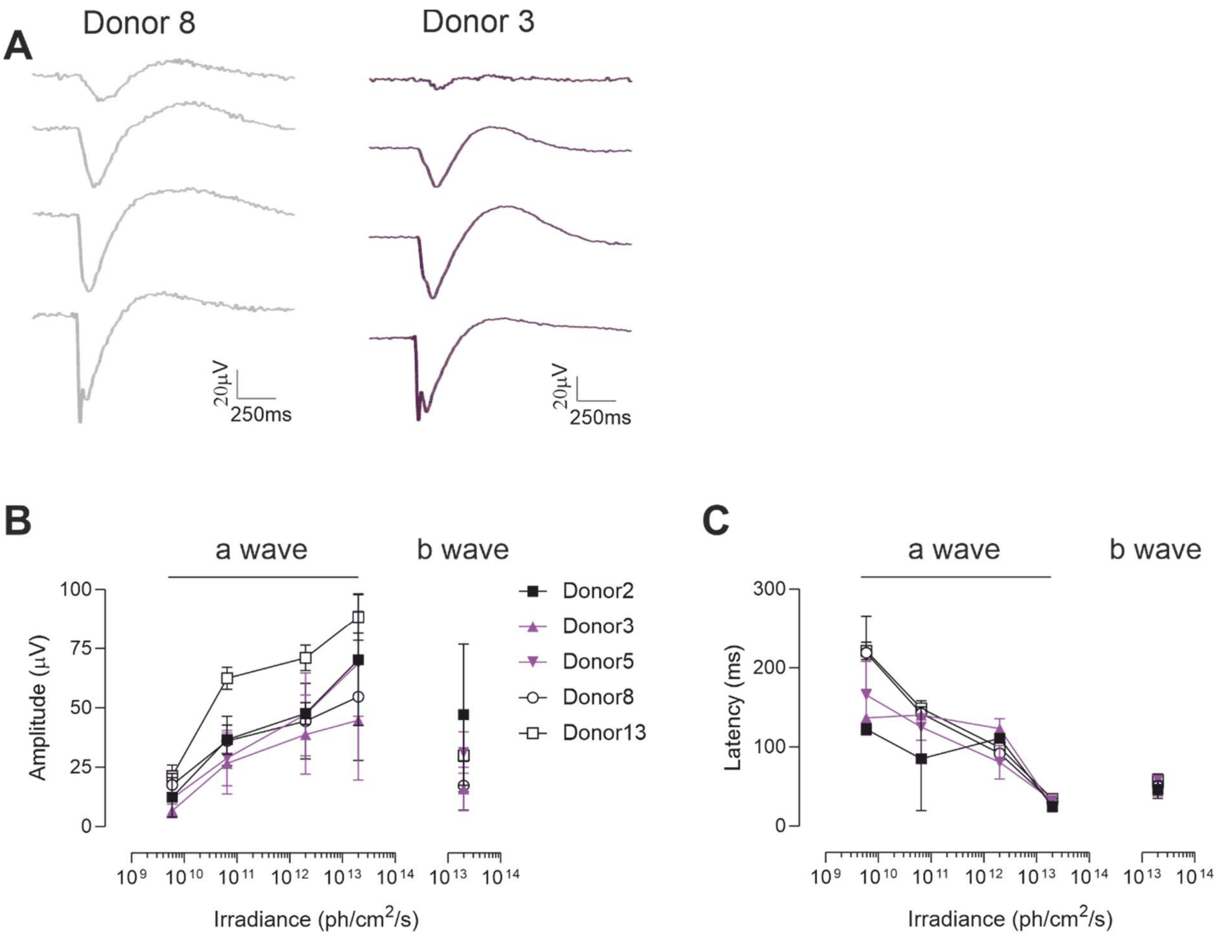


**Supplemental Figure 3: *ex vivo* micro Electroretinogram (mERG).** (**A**) Representative mERG traces from AD and age-matched donor in response to 10ms light pulses of increasing irradiance ($6.10^9$, $7.10^{10}$, $2.10^{12}$, and $2.10^{13}$ photons/cm$^2$/s, 470nm) (average of n=247 and n=244 mERG respectively). Dose response curves reporting the average amplitude (**B**) and latency (**C**) of the a and b waves in donors 2, 3, 5, 8, and 13 in response to 10ms light pulses of increasing irradiance ($6.10^9$, $7.10^{10}$, $2.10^{12}$, and $2.10^{13}$ photons/cm$^2$/s, 470nm) (n=2, 3, 3, 2, and 3 retina pieces in donors 2, 3, 5, 8, and 13 respectively).

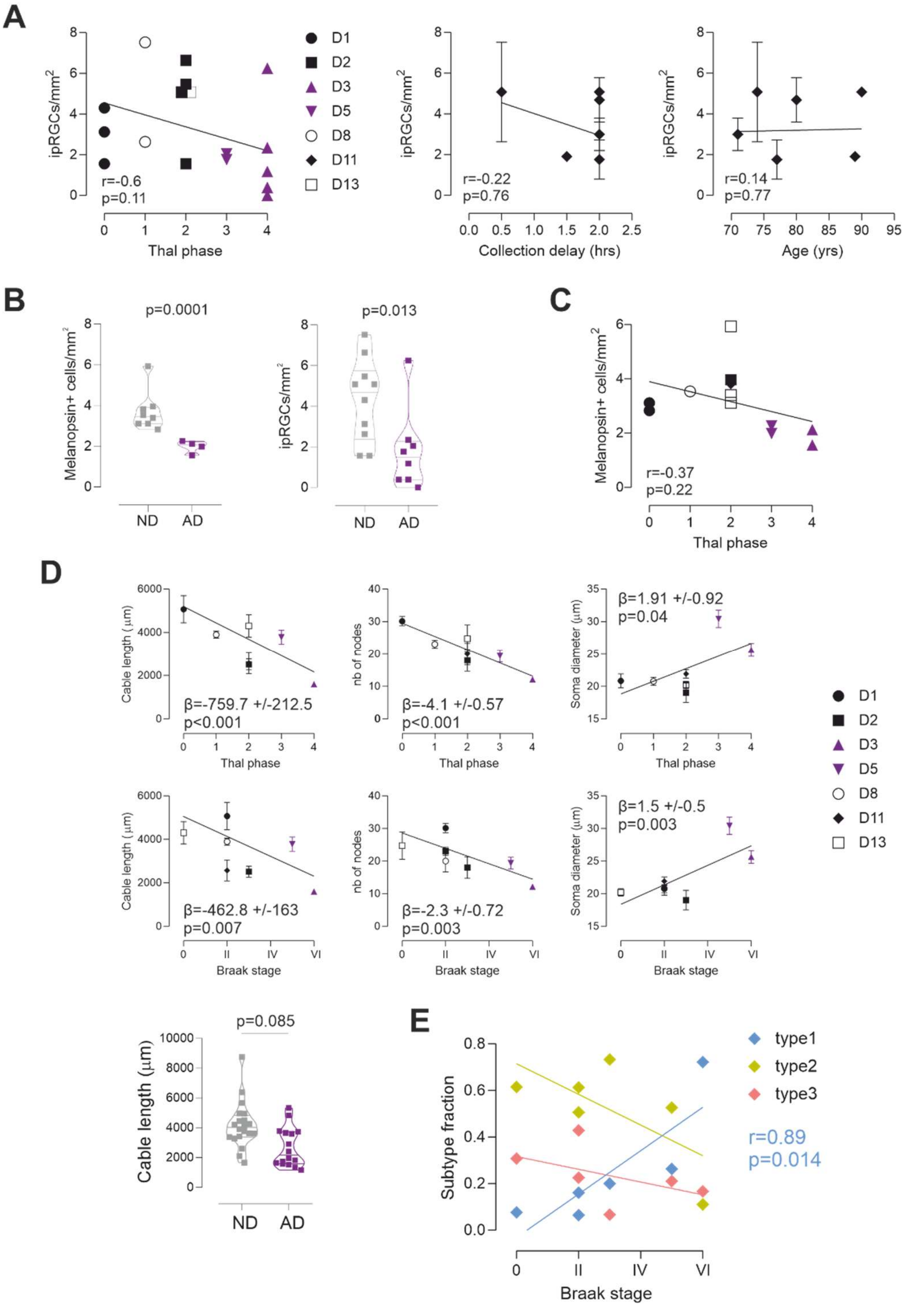


**Supplemental Figure 4: ipRGC abundancy and morphology decline in AD.** (**A**) ipRGC density do not correlate with either the Thal phase, age of the donor, or delay of eye collection. (**B**) Comparison of density of mel+ cells and ipRGCs measured in patches of retina from ND and AD donors (LMM, *left* $F(1,10)= 38.9$, *right* $F(1,16)=7.91$) . (**C**) Mel+ cell density in pieces of retina from each donor as a function of their Thal phase. (**D**) Correlation of each mel+ cell measurement (neurites length, number of nodes, and soma diameter) with Thal phases and Braak stages of the donors (*upper* and *lower* rows respectively) (LMM, slope β +/-SE) and cable length comparaison between AD (n=8 cells per donor, 2 donors, *purple*) and ND (n= 4 to 8 cells per donor, 4 donors, *grey*) (LMM, $F(1,38)=3.13$). (**E**) ipRGC subtypes distribution as a function of Braak stage.
